# Tube feet dynamics drive adaptation in sea star locomotion

**DOI:** 10.1101/2025.04.22.649740

**Authors:** Amandine Deridoux, Sina Heydari, Stanislas N. Gorb, Eva A. Kanso, Patrick Flammang, Sylvain Gabriele

## Abstract

Sea stars use hundreds of tube feet on their oral surface to crawl, climb, and navigate complex environments—all without the coordination of a central brain. While the morphology of tube feet and their role as muscular hydrostats are well described, the dynamics underlying their locomotion remain poorly understood. To investigate these dynamics, we employed an optical imaging method based on frustrated total internal reflection to visualize and quantify tube foot adhesion in real time across individuals of *Asterias rubens* spanning a wide size range. Our results reveal an inverse relationship between crawling speed and the duration of tube foot contact with the substrate. This suggests that sea stars regulate locomotion by modulating foot-substrate interaction time in response to body load. To test this, we conducted perturbation experiments using 3D-printed backpacks that increased body mass by 25% and 50%, along with numerical simulations based on a mechanistic model incorporating decentralized feedback control of the tube feet. The added load significantly increased adhesion time, supporting the role of a load-dependent mechanical adaptation. We further investigated inverted locomotion, both experimentally and through simulation, and found that tube feet adjust their contact behavior when the animal is oriented upside down relative to gravity. Together, our findings demonstrate that sea stars adapt their locomotion to changing mechanical demands by modulating tube foot-substrate interactions, revealing a robust, decentralized strategy for navigating diverse and challenging terrains.

## Introduction

Sea stars are commonly found in numerous marine environments, from tide pools to the deeper waters of the continental shelf, where they engage in diverse activities, including locomotion for foraging or posture maintenance during food consumption. These activities rely on their ability to maintain their oral surface in contact with the substratum (1), allowing them to navigate and adapt to their environments (2). A critical component enabling these functions is the coordinated action of tube feet which are specialized locomotory and adhesive appendages. Although tube feet have been well-characterized morphologically and mechanically (3–5), the way in which they collectively enable locomotion in sea stars remains poorly understood.

Tube feet consist of a flexible stem with an enlarged, flattened disc at the tip. Adhesion is mediated by the disc which allows strong yet temporary attachment to surfaces. The disc epidermis conforms to the substratum profile and secretes a proteinaceous adhesive material to secure the tube foot to the surface(6). This process is thought to involve a duo-gland adhesive system, in which one type of glandular cell secretes adhesive proteins, while another produces de-adhesive substances to enable detachment (7–9). In the species *Asterias rubens,* the chemical composition of these adhesive secretions has been studied extensively, providing insight into the mechanisms underlying their temporary adhesion (10, 11)

The stem, on the other hand, enables movement through a process powered by the water vascular system (12). This hydraulic system, filled with a fluid similar to seawater but with higher potassium concentration and osmolarity, generates the hydrostatic pressure necessary for tube foot operation ^9–11^. The stem of each tube foot, a hollow cylinder, provides both mobility and tensile strength (13). When internal pressure increases, the tube foot elongates, contacts the substrate, and adheres temporarily. The contraction of longitudinal muscle bundles then bends the tube foot. After detachment, muscle contraction shortens the stem, forcing the fluid back into the ampulla (1, 3). This coordinated muscle action enables the stepping movement of the tube feet (14). This movement is integral to sea star locomotion,. Indeed, despite lacking a centralized nervous system (15), sea stars are capable of highly coordinated locomotion, with crawling identified as their primary mode of movement (16–20).

Across animal species, locomotion speed typically scales with body size or mass, though this relationship often does not hold within a single species (21). In sea stars, locomotor capacity is expected to increase with body size, potentially due to a corresponding increase in the number and/or size of tube feet (22). Interestingly, this idea is somewhat counterintuitive: in many other animals, increased speed is often achieved by reducing the number of legs involved in locomotion, while a greater number of limbs is usually associated with generating stronger propulsion or enhancing adhesion. In sea stars, however, the relationship appears to vary significantly between species, underscoring the need for a more detailed understanding of tube foot dynamics (16, 22).

To improve our understanding of the role of tube feet in sea star locomotion, this study focuses on the dynamics of tube feet in *A. rubens*. In this species, tube feet are arranged in four rows on the oral side of each arm (2). Each individual therefore possesses hundreds of tube feet that must act cooperatively to enable crawling. Recognizing the central role of tube foot dynamics in sea star locomotion, we designed an innovative experimental approach to characterize these mechanisms in unprecedented detail. A major limitation of previous studies has been the inability to unambiguously identify adhering tube feet and their adhesion dynamics. To overcome this challenge, we implemented an innovative method based on frustrated total internal reflection (FTIR) imaging. This technique allowed us to quantify the number of adhering tube feet, and their attachment-detachment dynamics in an automated and precise manner, so that we could link these parameters to crawling speed and morphometry. Additionally, to support out experimental findings, we adapted a mechanistic model of the sea star with decentralized tube feet control (3) to investigate crawling under increased weight and in inverted locomotion, providing further insights into the adaptive biomechanical strategies of the movement.

## Results

### Allometric scaling in sea stars

We first investigated the scaling relationship between the body mass and the average arm length across eight sea stars species with five arms **(Fig. 1a)**. The log-log displays a strong positive correlation, suggesting a power-law (R^2^=0.8543) relationship between these two parameters. This result supports the hypothesis that arm length scales predictably with the body mass across species. Then, we focused on the morphology of sea stars of the species *A. rubens* **(Fig. 1b)**, our model species, across a broad size range represented by 24 specimens **(Fig. 1c)**. The characteristic pentaradial symmetry of *Asterias rubens* remains consistent over varying sizes. **Fig. 1c** further exemplifies this variability, showcasing the range of sizes analyzed in this study. To assess size-related parameters, the sea star average arm length was calculated as the mean of the distances from the central mouth to the tip of each of the five arms. A positive correlation was observed between average arm length and mass of individuals **(Fig. 1d)**, indicating that heavier sea stars are predictably larger, consistent with biological scaling principles.

**Figure 1.**
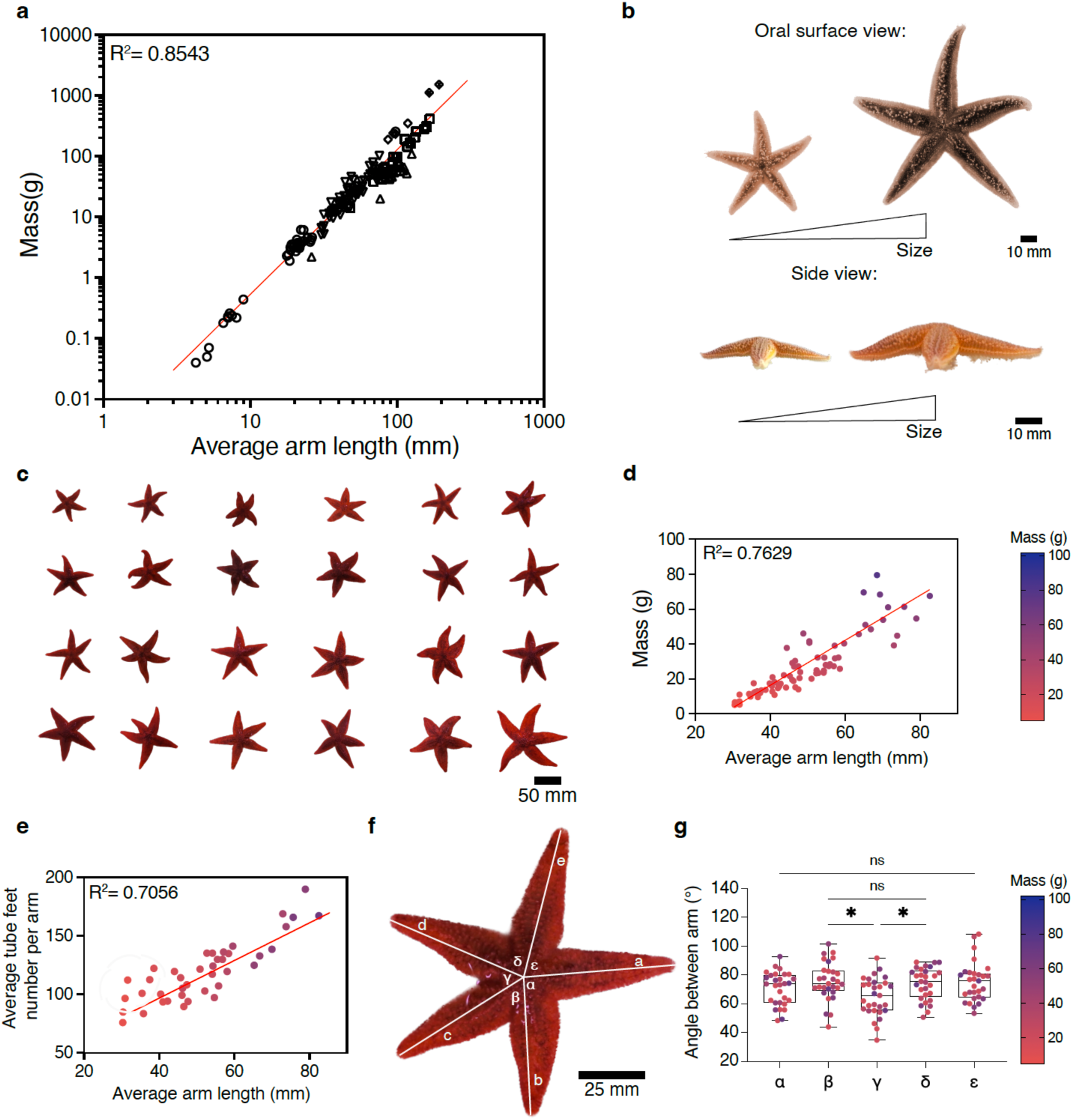
Allometric scaling in sea stars. **(a)** Relationship between the sea star mass and the average arm length across eight five-armed species: *Asterina gibossa* (circles, n = 37), *Echinaster sepositus* (triangles, n=14), *Asterias rubens* (inverted triangles, n=74)*, Marthasterias glacialis* (squares, n=23)*, Linckia laevigata* (squares with dot, n=2)*, Patiria miniata* (lozenges, *n = 4), Dermasterias imbricata* (lozenges with dot, n=2), and *Pentaceraster mammillatus* (circles with dot, n=1). Each dot represents an individual sea star (n = 157). The relationship shows a strong correlation (R^2^ = 0.8543, *p* < 0.0001). **(b)** Photographs of the oral and lateral views of two individuals of *Asterias rubens* illustrating size variability. **(c)** Size range observed in *A. rubens*. The scale bar is 50 mm. **(d)** Relationship between the body mass and the average arm length (*n* = 74, R^2^ = 0.7629, *p* < 0.0001). **(e)** Relationship between the average tube feet number per arm and the average arm length of *A. rubens* individuals (*n* = 40, R^2^=0.7056, *p* < 0.0001). **(f)** Aboral view of *A. rubens* showing the labeled assignment of arms and the corresponding inter-arm angles. The scale bar is 25 mm. **(g)** Measurements of inter-arm angle (*n* = 29). All data are color-coded by individual mass with *p < 0.05 and ns is not significant.

To confirm these results, we investigated morphology in another sea star species, *Marthasterias glacialis*, also across a broad size range **(Supplementary Fig. 1a)**. The same positive correlation between the average arm length and the body mass was observed **(Supplementary Fig. 1b)**. We also determined the average number of tube feet per arm across the full range of sea stars with varying masses and showed it increased linearly with size and mass (**Fig. 1e**). We next examined whether the pentaradial symmetry in *A. rubens* is maintained during locomotion by labeling the arms in a clockwise manner as a, b, c, d, and e (**Fig. 1f)**, using the leading arm “a” as a reference for analyzing angular arrangements. Interestingly, the angular analysis (**Fig. 1g**) revealed a subtle asymmetry: with the angle (γ) opposite to the leading arm “a” is significantly smaller (64.9±13.3°) compared to other inter-arm angles. This suggests that although pentaradial symmetry is a defining characteristic of *A. rubens,* minor deviations in arm angles occur and may be functionally associated with locomotion, independently of body size or mass.

### Quantification of the tube feet dynamics

Locomotion trials were conducted in an experimental glass aquarium, where seawater temperature was maintained at 15°C and salinity was set to 33‰. To ensure that glass was a suitable substrate for studying sea star locomotion, we performed detachment trials using a force ramp protocol to assess adhesion capabilities **(Supplementary Fig. 2a-b and Supplementary Movie 1)**. Additionally, we compared locomotion on glass with that on slate—a rougher, more naturalistic surface for sea stars **(Supplementary Fig. 2c).** Crawling speed did not differ significantly between glass 0.98±0.38 mm/s and slate 1.12±0.22 mm/s surfaces, confirming that glass is an appropriate model surface for studying sea star locomotion. To visualize the contact between the tube feet of *Asterias rubens* and the substrate, we used a custom-built aquarium made of high-refractive-index glass (n_1_=1.52) equipped with an LED strip at its base (**Fig. 2a-b and Supplementary Movie 2**). This setup leveraged the principle of frustrated total internal reflection. Under normal conditions, light traveling through the glass undergoes total internal reflection when it reaches the glass-seawater interface (n_2_=1.34) at an angle exceeding the critical value, effectively "trapping" the light within the glass (**Fig. 2a**). However, when a tube foot makes close contact with the glass, it alters the local refractive index, allowing light to escape and diffuse into the second medium. This interaction illuminates the contact area, producing bright dots visible at points of adhesion (**Fig. 2c and Supplementary Movie 3**).

**Figure 2.**
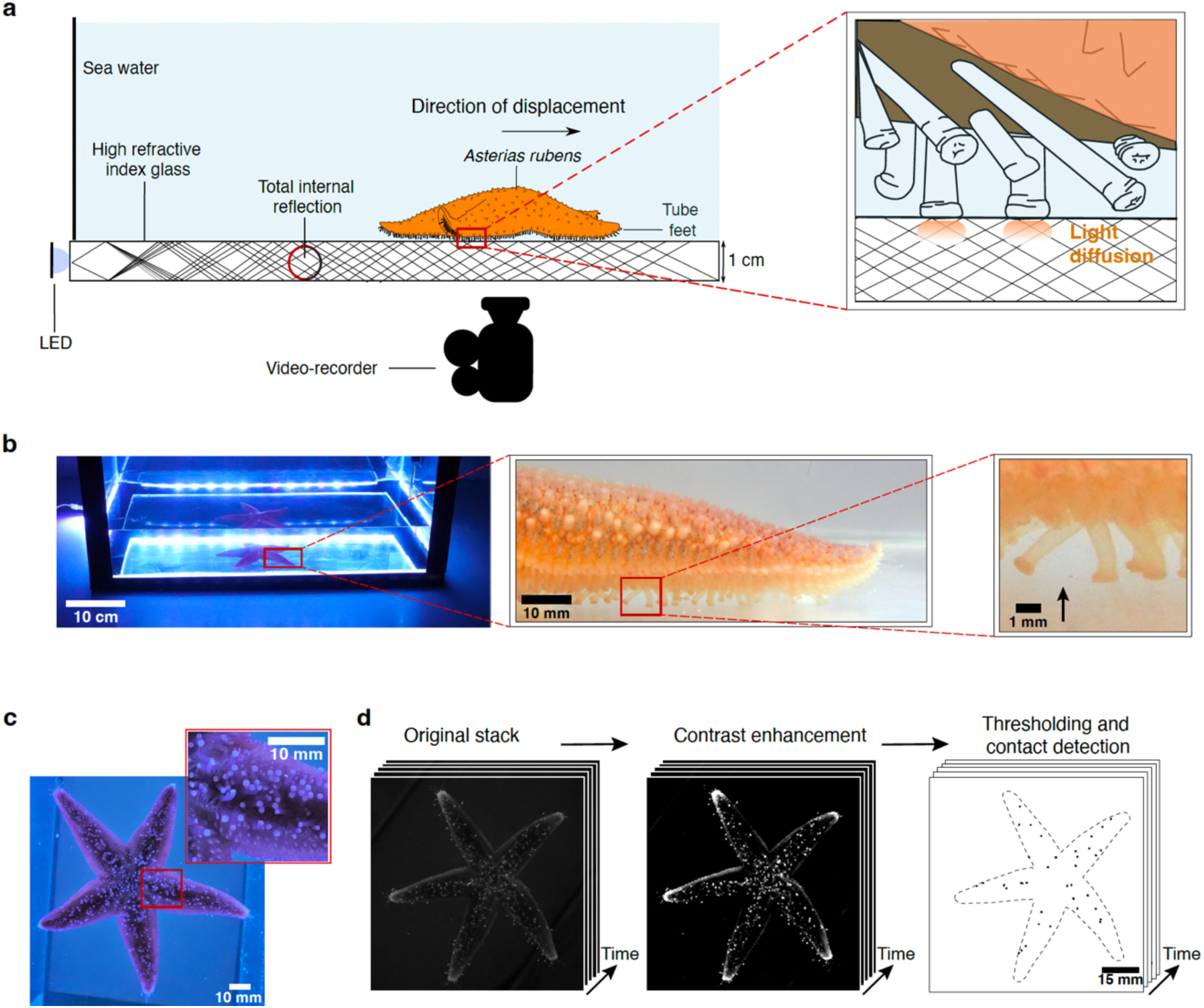
Quantification of tube foot dynamics using FTIR-based imaging. **(a)** Schematic representation of the experimental setup, which adapts the principle of frustrated total internal reflection (FTIR) to a custom-designed aquarium. When tube feet make contact with the FTIR-equipped glass surface, they disrupt the total internal reflection, causing light to scatter and locally illuminate the contact area. Sea stars are allowed to move freely, and their movements are recorded from below using a camera positioned beneath the tank. **(b)** Photograph of the experimental setup showing the aquarium equipped with a FTIR-based imaging system. The inset provides a close-up view of an *Asterias rubens* arm, where contact points of individual tube feet with the substrate are clearly visible during locomotion (arrow). **(c)** Image of the oral surface of *Asteria rubens* crawling within the experimental setup. **(d)** Image analysis pipeline developed to quantify the number of adhering tube feet and their contact area over time from image sequences. The method includes contrast enhancement, thresholding, and automated detection for accurate temporal tracking.

The oral surface of the sea star was filmed using a camera positioned beneath the aquarium, allowing for precise visualization and quantification of the tube foot adhesion during locomotion. Before focusing on the quantification of adhering podia, we determined the average number of tube feet per arm across the full range of sea stars with varying masses. To analyze the attachment-detachment dynamics, we developed a three-step image processing and thresholding method (**Fig. 2d and Supplementary Movie 4**). This method involves processing the video frames to enhance contrast, isolating the adhesion sites, and quantifying the contact area over time. This approach provided high-resolution insights into tube foot contact behavior, offering a robust platform to study the dynamics of attachment in sea star locomotion.

### Crawling speed in *Asterias rubens* is not correlated with the average number of tube feet in contact

The locomotion of individuals of *A. rubens* was recorded over 20 s, during which the number of tube feet in contact with the substrate was dynamically monitored **(Fig 3a)**. These trials revealed that the average number of tube feet in contact **(Fig. 3b)** remained relatively constant during locomotion, with only minor fluctuations observed over time.

**Figure 3.**
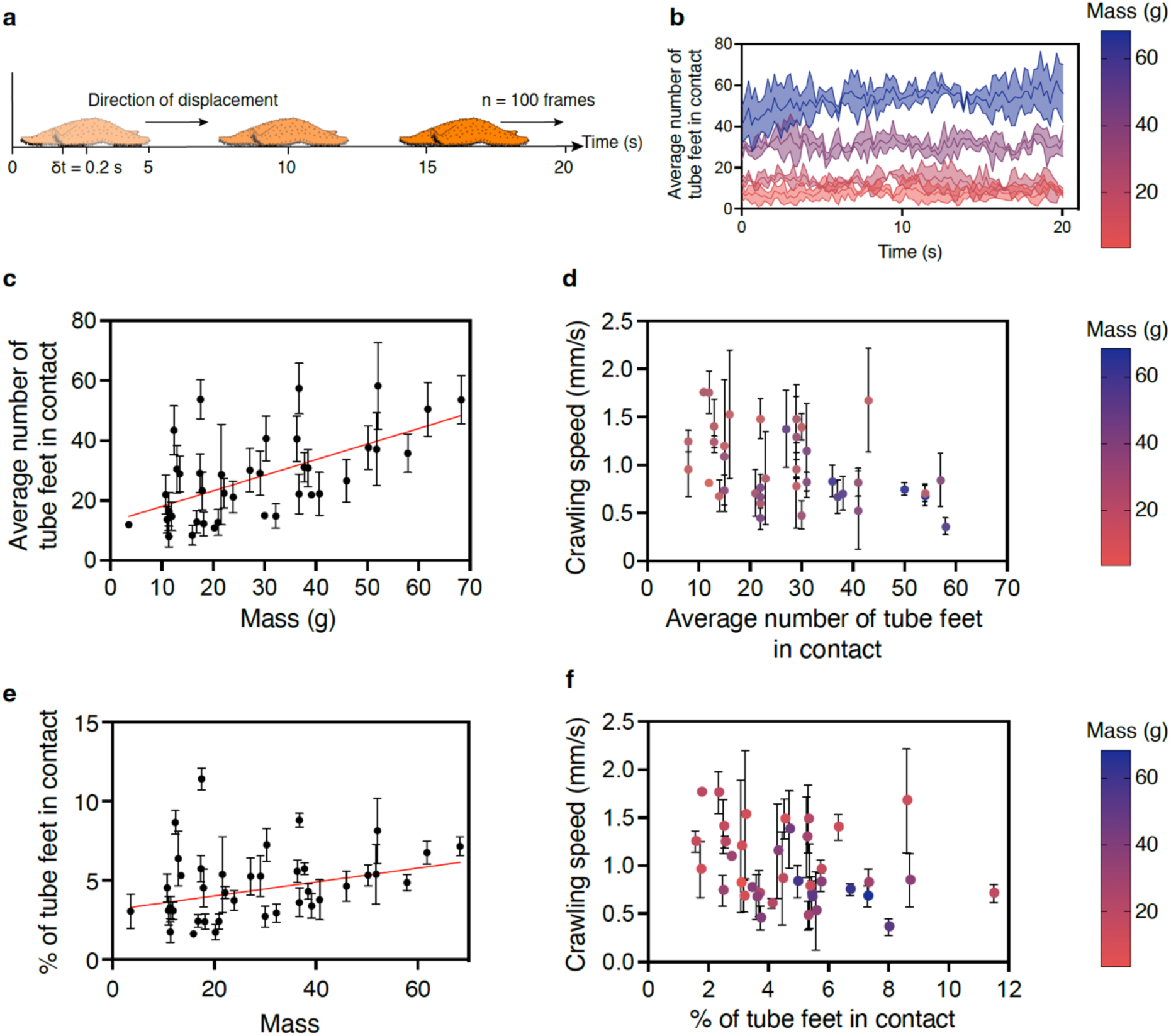
Crawling speed in *Asterias rubens* is not affected by the number of adhering tube feet. **(a)** Displacement of *A. rubens* recorded over 20 s, with the number of tube feet in contact and the contact area measured every 0.2 s. **(b)** Average number of tube feet in contact over time for individuals of increasing mass (color-coded from light to dark: 11.39 g, 20.96 g, 37.75 g, and 68.31 g). **(c)** Relationship between the average number of tube feet in contact and *A. rubens* mass (*n* = 39, R^2^ = 0.3536, *p* < 0.0001). **(d)** Crawling speed as a function of the average number of tube feet in contact and body mass (*n* = 39). **(e)** Relationship between the percentage of tube feet in contact and body mass (n = 39, R^2^ = 0.09304, *p* = 0.0002). **(f)** Relationship between the crawling speed and the average percentage of tube feet in contact (n = 39). All data are presented as mean ± standard deviation (s.d.).

A significant finding from these experiments is the linear relationship between the average number of tube feet in contact and natural body mass both for *Asterias rubens* **(Fig. 3c)** and *Marthasterias glacialis* **(Supplementary Fig. 3a)**. Larger individuals have an increased number of tube feet present on their oral surface, as the radius length scales with body size **(Fig. 1d)**. This relationship is consistent with established scaling principles, where morphological traits tend to increase with body mass (13). Interestingly, no significant relationship was found between crawling speed and the average number of tube feet in contact with the substrate in either *A. rubens* (**Fig. 3d)** or *M. glacialis* (**Supplementary Fig. 3b**), suggesting that sea stars maintain a relatively constant crawling speed regardless of how many tube feet are engaged. Despite having a large number of tube feet on their oral surface, only a small fraction—approximately 10%—is actively used during locomotion **(Fig. 3e and Supplementary Fig. 4a**), consistent with previous observations (23). Furthermore, crawling speed showed no correlation with the percentage of tube feet in contact **(Fig. 3f and Supplementary Fig. 4b)**. This apparent constancy may reflect compensatory mechanisms involving tube foot adhesion dynamics, supporting robust locomotion across individuals of varying sizes and species.

### Tube foot adhesion time drives adaptive and efficient sea star locomotion

To further investigate adhesion dynamics, high-magnification videos of the ambulacral groove and side views videos were recorded. These recordings provided detailed visualizations of tube foot dynamics, revealing three distinct stages **(Fig. 4a and Supplementary Movie 5)**. The locomotion cycle begins with the attachment stage, during which the tube foot approaches the substrate at an angle, resulting in an elliptical contact shape and a circularity index less than one. This is followed by the adhesion stage, where the tube foot remains firmly attached to the substrate—referred to as the adhesion time—characterized by a circularity index close to one, reflecting a near-perfect, vertical contact. Finally, in the detachment stage, the tube foot releases from the substrate in preparation for the next movement cycle. Temporal variations in the circularity index reveal therefore fluctuating patterns that allow precise determination of the period corresponding to the adhesion time of each individual tube foot **(Fig. 4b)**.

**Figure 4.**
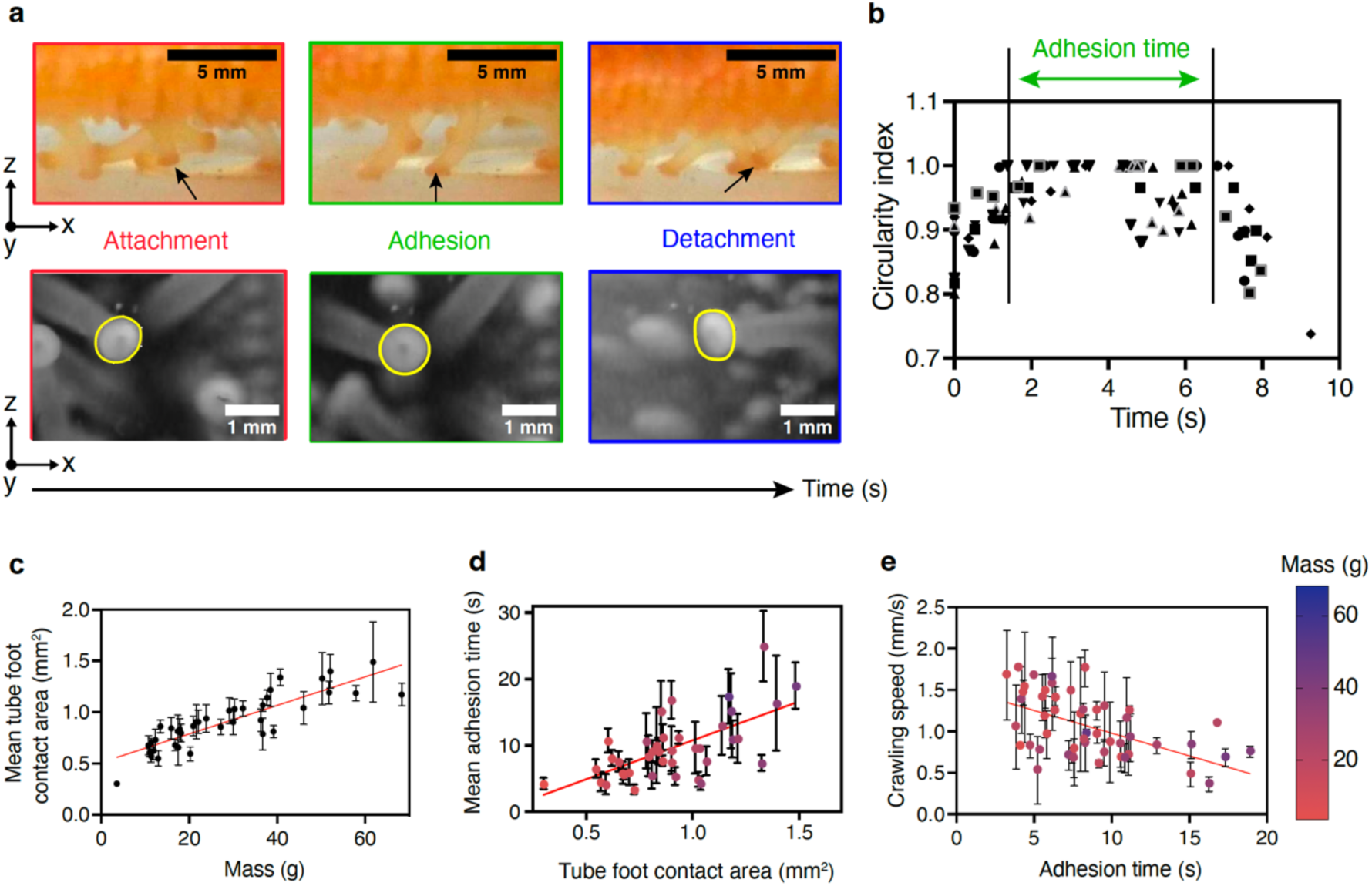
Tube foot adhesion time correlates with sea star locomotion in *Asterias rubens*. **(a)** High-magnification imaging reveals the three stages of tube foot adhesion during *Asterias rubens* locomotion. Top panels show side views of an individual in motion, while bottom panels display views of the ambulacral groove. Adhesion time is defined as the duration of the adhesion stage. **(b)** Temporal variation in the circularity index of individual tube feet (*n* = 10). Vertical lines indicate the start and end of the adhesion phase. **(c)** Relationship between mean tube foot contact area and body mass (*n* = 39; R² = 0.6203, *p* < 0.0001). **(d)** Relationship between mean adhesion time and tube foot contact area (*n* = 39; R² = 0.3088, *p* < 0.0001). **(e)** Crawling speed as a function of tube foot adhesion time (*n* = 49; R² = 0.2115, *p* < 0.0001). All data are presented as mean ± standard deviation (s.d.).

Analysis of the mean tube foot contact area during the adhesion stage shows a positive correlation with natural body mass **(Fig. 4c and Supplementary Fig. 5a)**. Larger sea stars exhibit allometrically scaled tube feet and contact areas, resulting in enhanced adhesive capabilities as predicted by scaling laws relating mass and surface area. Additionally, a weak but significant positive relationship was observed between contact time and contact area **(Fig. 4d)**, indicating that larger tube feet require slightly longer contact times.

We also found an inverse relationship between crawling speed and contact time **(Fig. 4e and Supplementary Fig. 5b)** demonstrating that longer contact times are associated with slower locomotion. This indicates that rapid detachment of tube feet is essential for achieving higher crawling speeds. Notably, contact times during locomotion remained below one minute, typically ranging from 3 to 20 s. Moreover, some tube feet appear to be reused during locomotion via repetitive detachment and attachment cycles, and the time between two successive adhesion stage is referred to as the recovery stroke phase (**Supplementary Fig. 6a**). This parameter remained relatively constant across individuals of different natural body masses, with an average duration of 18.6 ± 6.5 s (**Supplementary Fig. 6b**).

Taken together, these findings suggest that the relationship between body mass and crawling speed is mediated through adhesion dynamics. They highlight the pivotal role of tube feet coordination and timing in enabling adaptive and efficient locomotion in *Asterias rubens*.

### Dynamic adjustments in tube foot contact time in response to mass changes

To validate these findings, we conducted perturbation experiments involving artificial mass alteration and inverted locomotion in *A. rubens.* In the first experiment, individuals were fitted with a 3D-printed backpack carrying additional weights equivalent to 25% or 50% of their body mass **(Fig. 5a and Supplementary Movie 6)**. The percentage of tube feet in contact with the substrate remained consistent across conditions (normal locomotion, empty backpacks, +25% mass, and +50% mass), indicating that *A. rubens* maintains a stable level of tube foot engagement regardless of additional load **(Fig. 5b)**. This suggests a compensatory strategy to preserve locomotor stability under increased mechanical demand.

**Figure 5.**
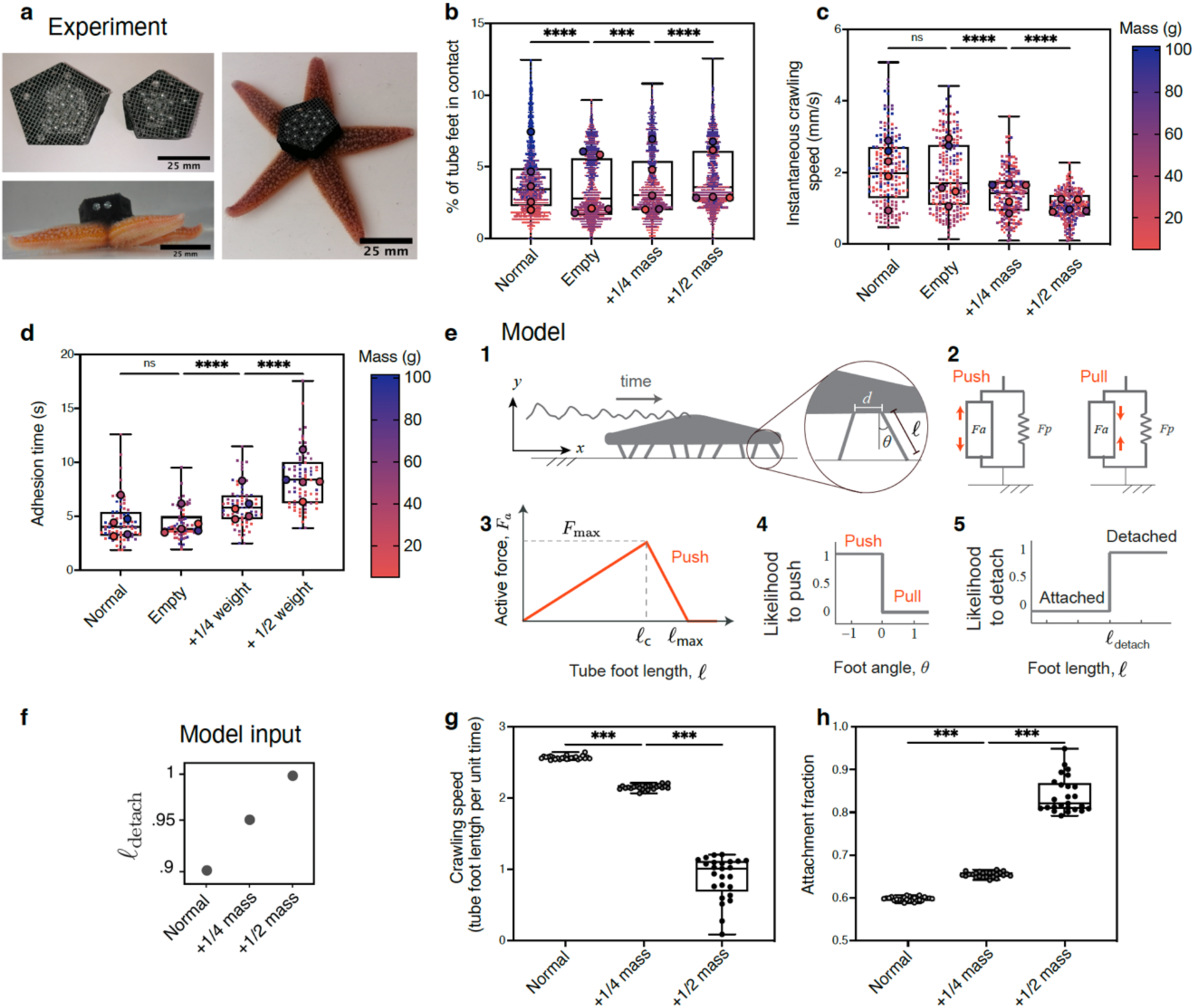
Sea star locomotion dynamics in response to changes in mass are modulated by adhesion time. **(a)** Photograph showing two representative sizes of 3D-printed backpacks used to artificially increase the mass of *Asterias rubens* by adding stainless steel beads. **(b)** Percentage of adhering tube feet, **(c)** instantaneous crawling speed, and **(d)** tube foot adhesion time across four conditions: normal locomotion, empty backpack, +25% body mass, and +50% body mass (N = 5 sea stars per condition). Each small dot corresponds to one tube foot contact measurement taken every 0.2 s. for the same four conditions. Data in all superplots are color-coded by mass; large dots indicate individual averages. Three replicates were performed per sea star. **(e)** Schematic of the mathematical model of sea star locomotion. (1) Model organism propelled by 10 tube feet. The inset defines key parameters: tube foot length (*l*), tilt angle (*θ*), and inter-foot spacing (*d*). (2) Mechanical model of a single tube foot, composed of a passive linear spring and an active force generator capable of producing either pushing or pulling forces depending on muscle activation. (3) Active force profile based on Hill’s muscle model. (4) Local control policies at the level of individual tube feet. These define the probability of transitions between pushing and pulling states and between attached and detached states, based on sensory feedback. Tube feet detach when stretched beyond a threshold length *l_detach*. **(g)** Simulated crawling speed and **(h)** attachment fraction τ, with 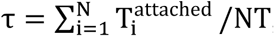, defined as the sum over all tube feet of the time 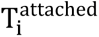 each foot spends in attachment, divided by the total simulation time T times the number N of tube feet. Both metrics are shown as a function of sea star weight, increasing from baseline to highest (*W* = 2, 2.5, 3). For each weight condition, 25 simulations were performed with randomized initial foot states. ***p < 0.001, ****p < 0.0001, ns = not significant.

Despite the constant tube foot engagement, instantaneous crawling speed declined significantly with added weight **(Fig. 5c**), likely due to the greater tube foot force required for tube foot detachment and propulsion. Correspondingly, adhesion time increased with mass, from 4.3±1.4s for the empty backpack, to 5.9±1.9s and 8.4±2.6s for +25% and +50% mass, respectively **(Fig. 5d)**. This prolongation likely reflects the increased propulsive force required to move the added weight during each locomotor cycle. Together, these results confirm that crawling speed is inversely related to contact time, which itself is modulated by body mass. They further highlight the adaptive control of tube foot adhesion dynamics as a key mechanism for maintaining effective locomotion under varying mechanical loads.

### Biomechanics model

We hypothesized that the observed increase in contact time with increasing body mass is governed by the biomechanics of the tube feet. To corroborate this hypothesis, we employed a mathematical model of sea star locomotion driven by tube feet forces (3). In this model, the sea star is represented as a rigid body of mass *m*, whose center of mass is located at (*x*, *y*) in the vertical plane. The ventral surface is lined by N tube feet, anchored at evenly spaced distance intervals *d* along the sea star body (**Fig. 5e)**. Each tube foot is characterized by its length ℓ and inclination angle *θ* from the vertical axis. When attached and engaged with the substrate, a tube foot exerts an active force *F*_*a*_, caused by the activation of muscle tissues lining the stem and ampulla, and experiences a passive restorative force *F*_"_due to connective tissue resistance. Both active and passive forces act along the length of the tube foot and can either push or pull on the sea star’s body, depending on which muscle group is activated: ampullar muscles contraction extends the tube foot and produce a pushing force, while stem contraction shorten the foot, resulting in a pulling force (3, 5).

We imposed no direct control on the sea star center of mass and considered a fully decentralized adaptive control of the tube feet. Each attached tube foot senses its own length and inclination angle (ℓ, *θ*) and responds accordingly. Depending on the sign of *θ*, the tube foot experiences shear either in the same or opposite direction to its horizontal motion and decides to either push or pull, following a Hill’s muscle model (**Fig. 5e**). When the tube foot is axially stretched beyond a critical length ℓ_#$%&’(_, it detaches. In the detached state, the tube foot applies no force and remains inactive for a stochastic time duration governed by a reattachment rate *λ*; the longer it remains detached, the higher its probability of reattaching. Upon reattachment, the tube foot makes a random step in the direction of the sea star’s motion. A full mathematical description of the model is provided in the Theoretical Supplementary Information. In the model, the characteristic length scale is set by the tube foot length, while the time scale is determined by the passive relaxation time of a foot returning to its resting length after deformation.

We examined numerically the sea star locomotion, while keeping all parameters the same, except for the sea star weight and detachment length ℓ_#$%&’(_: namely, we increased the sea star weight by 25% and 50%, consistent with our experiments, and correspondingly, increased ℓ_#$%&’(_ with increasing the sea star weight (**Fig. 5f and Supplementary Movie 7**). For each weight, we performed 25 Monte-Carlo simulations, corresponding to random initial conditions of the tube feet. We calculated the crawling speed, defined as the total distance traveled by the center of mass divided by the total simulation time *T*. We also calculated the attachment fraction, 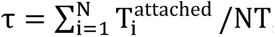, defined as the sum over all tube feet of the total time each foot spends in attachment 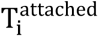, divided by *NT*. This gives the average proportion of time that a typical tube foot is attached. Interestingly, theoretical simulations of crawling speed **(Fig. 5g)** and attachment fraction **(Fig. 5h)** in response to mass variation show trends similar to those observed experimentally **(Figs. 5c–d)**, supporting the validity of our model based on fully decentralized adaptive control of the tube feet. These findings support the hypothesis that sea star locomotion can emerge from local sensing and mechanical feedback at the level of individual tube feet, without requiring centralized control. Moreover, they demonstrate that crawling speed under increased mechanical load can be modulated through local adaptation of detachment timing.

### Dynamic adjustments in tube foot adhesion time in inverted setup

In the second perturbation experiment, we analyzed inverted locomotion by placing individual’s upside down **(Fig. 6a and Supplementary Movie 8),** a condition that mimics the natural challenge of navigating vertical or overhanging surfaces in their environment. This inversion alters the distribution of gravitational forces, effectively increasing the load on the tube feet as they counteract the downward pull, thereby requiring greater adhesive forces and coordination to maintain stability and locomotion. The mean percentage of tube feet in contact slightly decreased in the inverted position compared to the normal condition **(Fig. 6b)**, suggesting a compensatory response to counteract the effect of gravity and maintain stability. Interestingly, instantaneous crawling speed was significantly reduced in the inverted condition **(Fig. 6c),** likely due to the higher energy demands associated with this altered posture. Adhesion time was also significantly prolonged during inverted locomotion **(Fig. 6d)**, consistent with the need for stronger adhesion to prevent detachment under the influence of gravity.

**Figure 6.**
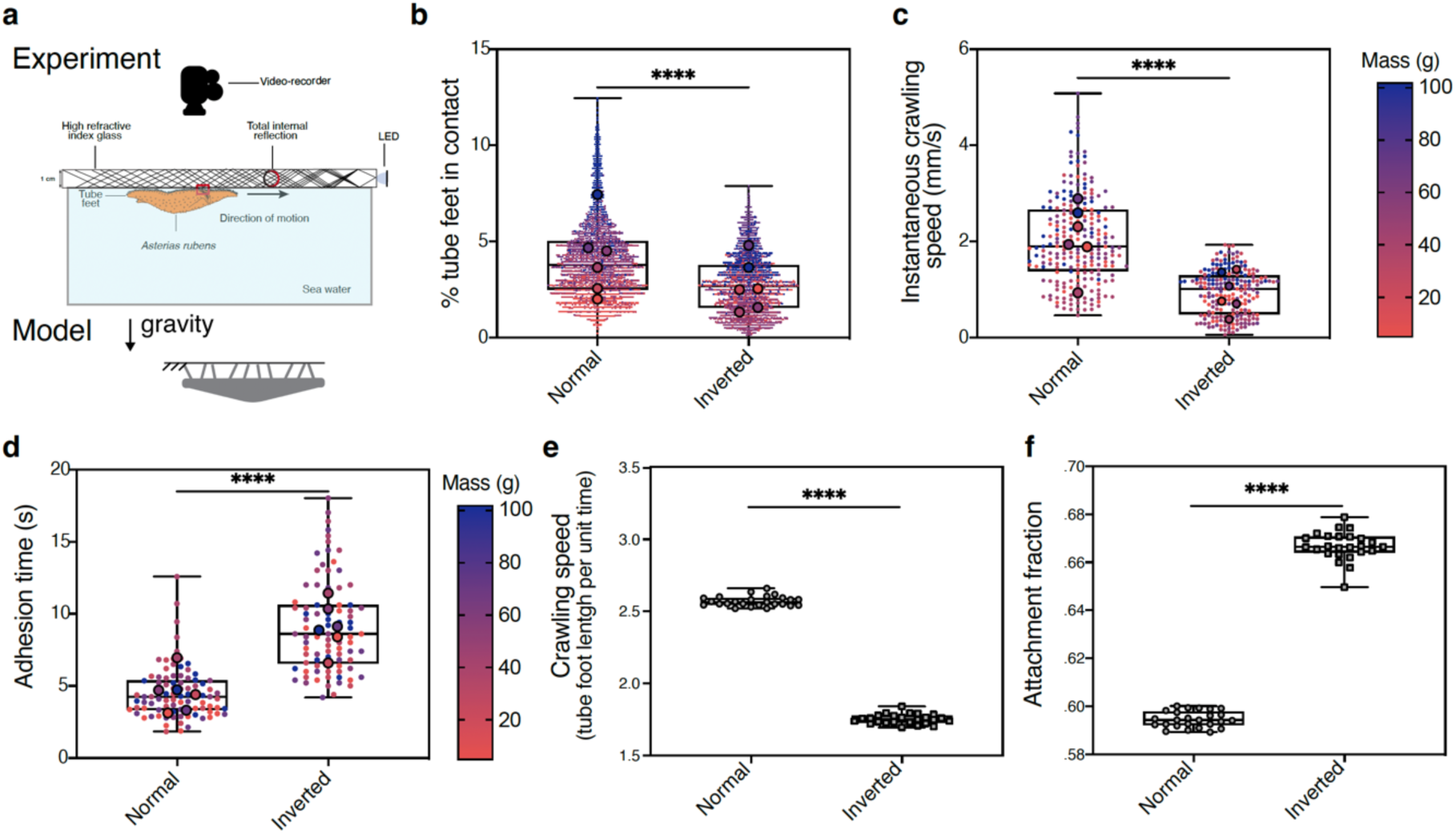
Modulation of adhesion time enables sea star to adapt their locomotion under inverted conditions. **(a)** Schematic of the experimental setup used to assess inverted locomotion. Sea stars were allowed to crawl upside down, with their oral surface recorded from above using a camera positioned over the aquarium. **(b)** Percentage of tube feet in contact, **(c)** instantaneous crawling speed, and **(d)** tube foot adhesion time under normal and inverted conditions for *Asterias rubens* under normal and inverted locomotion (*N* = 6 sea stars per condition). All data are displayed as superplots, showing individual data points (small dots representing an individual contact event) and sea star means (large dots). Each individual was tested in three replicates. **(e)** Simulated crawling speed and **(f)** attachment fraction 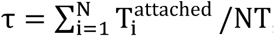, defined as the sum over all tube feet of the time 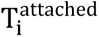 each foot spends in attachment, divided by the total simulation time T times the number N of tube feet. Both shown for the flat and inverted sea stars. For each orientation 25 random simulations have been performed. Parameter values are N = 100, L = 40, l = 1, F_max_ = 0.4, λ = 5, with no parameter adjustment between normal and inverted simulations. All data are displayed as superplots, showing individual data points (small dots) and sea star means (large dots). Each individual was tested in three replicates, and ****p < 0.0001.

We compared numerically, in the context of our model, the sea star locomotion under normal and inverted conditions, with no adjustment of parameters, keeping all parameters constant **(Supplementary Movie 9)**. Again, for each condition, we performed 25 Monte-Carlo simulations, corresponding to random initial conditions of the tube feet, and calculated the crawling speed and attachment fraction, 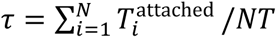. The simulation results show trends similar to those observed experimentally **(Fig. 6e–f)**, confirming the validity of the biomechanical approach.

Together, these perturbation experiments demonstrate that sea stars employ dynamic adjustments in tube foot adhesion time to adapt to increased mechanical demands, whether due to added weight or altered locomotion modes. These experimental findings, combined with the modeling results, underscore the critical role of tube foot dynamics in stabilizing and adapting sea star locomotion under varying physical constraints.

## Discussion

While the biomechanics of tube feet has been described in echinoderms, the dynamics of their contact formation and breakage during locomotion and the number of tube feet involved in movement have remained largely unexplored. Unlike many animals, sea stars display a less straightforward relationship between body mass and crawling speed (16, 20, 22, 24), prompting the need for a deeper examination of the mechanisms underlying their locomotion, especially by focusing on the role of their tube feet (16).

Our study demonstrates that *Asterias rubens* maintains pentaradial symmetry across a wide size range. The observed morphological consistency, along with only minimal angular deviations between arms, likely reflects evolutionary pressures to preserve efficient, symmetrical locomotion (3),(4). This robust geometry may support coordinated activation of tube feet and enhance mechanical stability during movement.

Although environmental factors, such as temperature, flow conditions, and substrate inclination are known to influence echinoderm locomotion (16, 22, 24, 25), our results show that the average crawling speed of *Asterias rubens* (0.98 ± 0.38 mm/s) is consistent with previous studies(26), validating our experimental design. Surprisingly, we found that crawling speed is not correlated with the average number or percentage of tube feet in contact with the substrate. Even though larger individuals naturally possess more tube feet, the proportion actively engaged in adhesion remains stable—suggesting that *A. rubens* utilizes a constant fraction of its podia for locomotion, independent of size. The lack of dependence on contact area further underscores an evolutionary adaptation that enables sea stars to maintain consistent and efficient locomotion across a variety of surfaces and environmental conditions. For instance, in *Acanthaster solaris*, locomotion speed is influenced by substrate texture, with slower movement observed on rougher surfaces (24). In *Patiria miniata*, some individuals possess five or six arms, yet the number of arms has been shown to have no significant effect on crawling speed (16). Additionally, comparisons across species indicate that sea stars inhabiting tropical environments generally exhibit faster locomotion than those from temperate regions (16).

Our observation highlights the importance of tube foot adhesion timing over mere contact number. Video recordings with a high pixel count revealed that locomotion consists of a cyclic sequence of attachment, adhesion, and detachment phases. While larger tube feet and greater contact areas likely enhance adhesion strength, they also lead to increased adhesion time, which in turn reduces crawling speed. These findings reveal a trade-off between adhesive performance and locomotor efficiency: rapid movement requires shorter adhesion times and thus faster detachment–reattachment cycles (27),(28). A positive correlation between stride frequency and maximum speed has been described in several animals using dynamic adhesion for locomotion (28).

Perturbation experiments show that *A. rubens* dynamically modulates tube foot contact time in response to added load. Although the number of adhering tube feet remained constant, crawling speed decreased and contact time increased proportionally with the added weight. This suggests that sea stars rely on local mechanical feedback to adaptively tune tube foot dynamics, without requiring changes in coordination or overall foot engagement. Notably, these same patterns were observed under inverted locomotion, where gravitational forces were altered but the percentage of tube feet in contact remained stable. The similarity of behavioral responses across both perturbations—additional mass and inversion—indicates a conserved mechanical strategy to stabilize movement under changing physical conditions.

Numerical simulations based on a biomechanical model of tube foot function replicated the observed trends. By varying body mass and detachment thresholds, the model reproduced the inverse relationship between crawling speed and adhesion time. The simulations further demonstrated that local sensing and adaptive control at the level of individual tube feet are sufficient to generate coherent and efficient crawling behavior—without centralized control. However, a centralized coordination in stepping direction is necessary for an efficient locomotion (23).

Together, our findings identify tube foot contact time (presumably correlating with adhesion time) as a key mass-dependent parameter that governs sea star locomotion. This mechanism provides a flexible strategy to balance stability and speed across individuals and environmental conditions. FTIR-based imaging has been previously developed to measure contact areas of adhesive pads during locomotion, but it has mostly been used for studying tree frog climbing, and stress distribution in gecko toes (29–31). Our FTIR platform offers a promising avenue for investigating locomotion dynamics in echinoderms and other soft-bodied aquatic animals.

## Conclusion

This study demonstrates that *Asterias rubens* modulates tube foot adhesion time to maintain effective locomotion across a range of body sizes and mechanical challenges. By combining high-resolution imaging, perturbation experiments, and mathematical modeling, we show that adhesion time—rather than the number of contacting tube feet—is the principal determinant of crawling speed. These results reveal a decentralized, mechanically adaptive strategy that enables robust and versatile locomotion in sea stars.

## Materials & Methods

### Collection and maintenance of sea stars

Specimens of *Asterias rubens* (Linnaeus, 1758) were hand-collected at low tide on breakwaters at Knokke-Heist, Belgium. Specimens of *Marthasterias glacialis* (Linnaeus, 1758) were hand-collected at the Concarneau marine station, France during low tide. They were brought back to the University of Mons, where they were housed in recirculating seawater aquaria maintained at a temperature of 12-13°C and a salinity of 30-33‰. Salinity was controlled using a precision salinity refractometer (ATC-S/Mill-E, ±0.2%). The sea stars were fed weekly with fresh mussels (*Mytilus edulis* L.). The collected individuals ranged in size from 60 mm to 180 mm in total diameter **(Fig. 1b)**. To minimize stress and allow for acclimatization, all individuals were kept under these conditions for at least two weeks prior conducting locomotion trials. Some of the specimens used in the experiments shown in **Fig. 1a** were kindly provided by the aquariology department of Nausicaá (Boulogne-sur-Mer, France).

### Morphometric analysis

To assess the size and morphological parameters of *Asterias rubens*, high-resolution images of the oral surface were taken in seawater before each experiment using Nikon 1 J5 camera equipped with a 1 Nikkor 10-30 VR lens. Videos and images were analyzed using Fiji(32). To measure the mass of each sea star, the oral surface was gently dried with a paper towel, and the specimen was weighed on a digital balance inside of a beaker full of sea water. To determine the number of tube feet per arm, individuals were anaesthetized in a 3.5% solution of magnesium chloride in seawater for ten minutes to relax their tube feet. Then, three arms were randomly selected, and the number of tube feet was counted to provide representative data on tube foot distribution.

### Measurements for locomotion experiments

The oral surface of *A. rubens* was video recorded using a Nikon 1 J5 camera with a 1 Nikkor 10-30 VR lens positioned beneath the aquarium. Each sea star was carefully placed at the bottom center of the aquarium, and its movement was recorded for 20 s once it began crawling. Measurements for locomotion assays included locomotion speed, the number of tube feet in contact with the substrate and the contact area during the locomotion. The percentage of tube feet in contact during locomotion was calculated for each individual using the following formula:

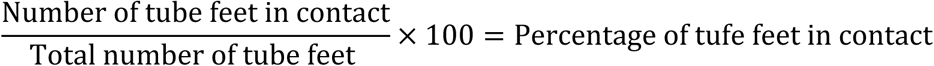

The instantaneous locomotion speed was calculated by dividing the displacement by the time required to move between two points. The displacement D between two successive positions (*x_1_, y_1_*) and (*x_2_, y_2_*) was determined using the following equation:

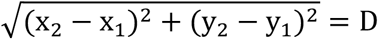

The dynamics of tube foot adhesion was analyzed by recording high-magnification videos of the oral surface of *A. rubens* during locomotion. These videos were captured using a Nikon D90 camera equipped with an AF-S Nikkor 105 mm lens. For these recordings, tube feet adhesion time and contact area were precisely measured, providing detailed insights into adhesion dynamics.

### Perturbation experiments

Backpacks were 3D-printed using an Ultimaker printer with PLA filament. The backpacks were securely attached to the sea stars using veterinary glue (Vetbond, 3M Deutschland GmbH) (**Fig. 5a)**. Locomotion was recorded under three conditions: with an empty backpack, and with additional weights equivalent to 25% and 50% of the sea star’s body mass. For each condition, movements were recorded for 20 s under the same setup as the experiments without backpacks. Stainless-steel marbles were used to provide the additional mass. In the inverted locomotion experiments, individuals of *A. rubens* were allowed to walk upside down on a high refractive index glass plate equipped with an FTIR-based system **(Fig. 6a)**. Locomotion was recorded using a Nikon 1 J5 camera with a 1 Nikkor 10-30 VR lens positioned above the plate to visualize their altered movement during inverted locomotion.

### Statistical analysis

Each experiment was repeated three times on each sea stars. Every set of data was tested for normality test using the D’Agostino Pearson test in Prism 10.0 (GraphPad) that combines skewness and kurtosis tests to test, whether the shape of the data distribution was similar to the shape of a normal distribution. For paired comparisons, significances were calculated in Prism 10.0 (GraphPad Software, Inc.) with Mann–Whitney (two-tailed, unequal variances). For multiple comparisons with non-normal distribution, data sets were analyzed with a Kruskal-Wallis test in Prism 10.0 (GraphPad Software, Inc.), which is a suitable nonparametric test for comparing multiple independent groups when the data are skewed. Linear regressions were used to analyze allometric scaling, to assess the relationship between average arm length and body mass and to investigate the relationship between crawling speed and various parameters. Unless otherwise stated, all data are presented as mean ± standard deviation (s.d.). The confidence interval in all experiments was 95% and as a detailed description of statistical parameters it is included in all figure captions with *p < 0.05, **p < 0.01, ***p < 0.001, ****p<0.0001 and n.s. is not significant.

## Acknowledgments

A.D., P.F. and S.G. acknowledge funding from the University of Mons ‒ UMONS, the Research Institute for Biosciences Project Starlight, FEDER Prostem Research Project no. 1510614 (Wallonia DG06), the F.R.S.-FNRS Epiforce Project no. T.0092.21, the F.R.S.-FNRS Project no. T.0088.20, the F.R.S.-FNRS Cellsqueezer Project no. J.0061.23, the F.R.S.-FNRS Optopattern Project no. U.NO26.22 and the Interreg projects MICROPLAITE and ANTIRESI, which are financially supported by Interreg France-Wallonie-Vlaanderen (Fonds Européen de Développement Régional, FEDER-ERDF). A.D. is financially supported by F.R.S.-FNRS as FRIA Grantee. S.G. acknowledges le Fonds pour la Recherche Médicale dans le Hainaut (FRMH). P.F. is Research Director of the F.R.S.-FNRS. A.D. and S.G. thank Stéphane Hénard from the Aquariology Department of Nausicaá (Boulogne-sur-Mer, France) for kindly providing access to various sea star species.

## Author contributions

S.G. and P.F. conceived the project and S.G. supervised the project. A.D. performed all experiments with sea stars and analyses. S.H. and E.A.K. developed the theoretical model and performed simulations and analysis. A.D., P.F. and S.G. analyzed data and prepared the figures with S.H. and E.A.K. The article was written by A.D. and S.G, read and corrected by all authors, who all contributed to the interpretation of the results.

