## Supplementary Information for "Tube feet dynamics drive adaptation in sea star locomotion"

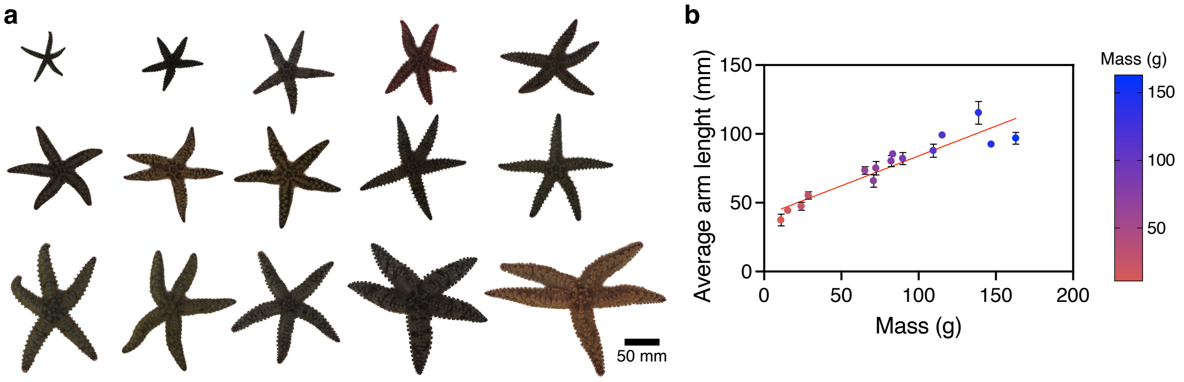


**Supplementary Figure 1 – Allometric scaling in *Marthasterias glacialis*. (a)** Size range investigated in *M. glacialis*. Scale bar: 50 mm. **(b)** Relationship between average arm length and body mass ($n = 15$; $R^2 = 0.8492$, $p < 0.0001$). All data are presented as mean ± standard deviation (s.d.).

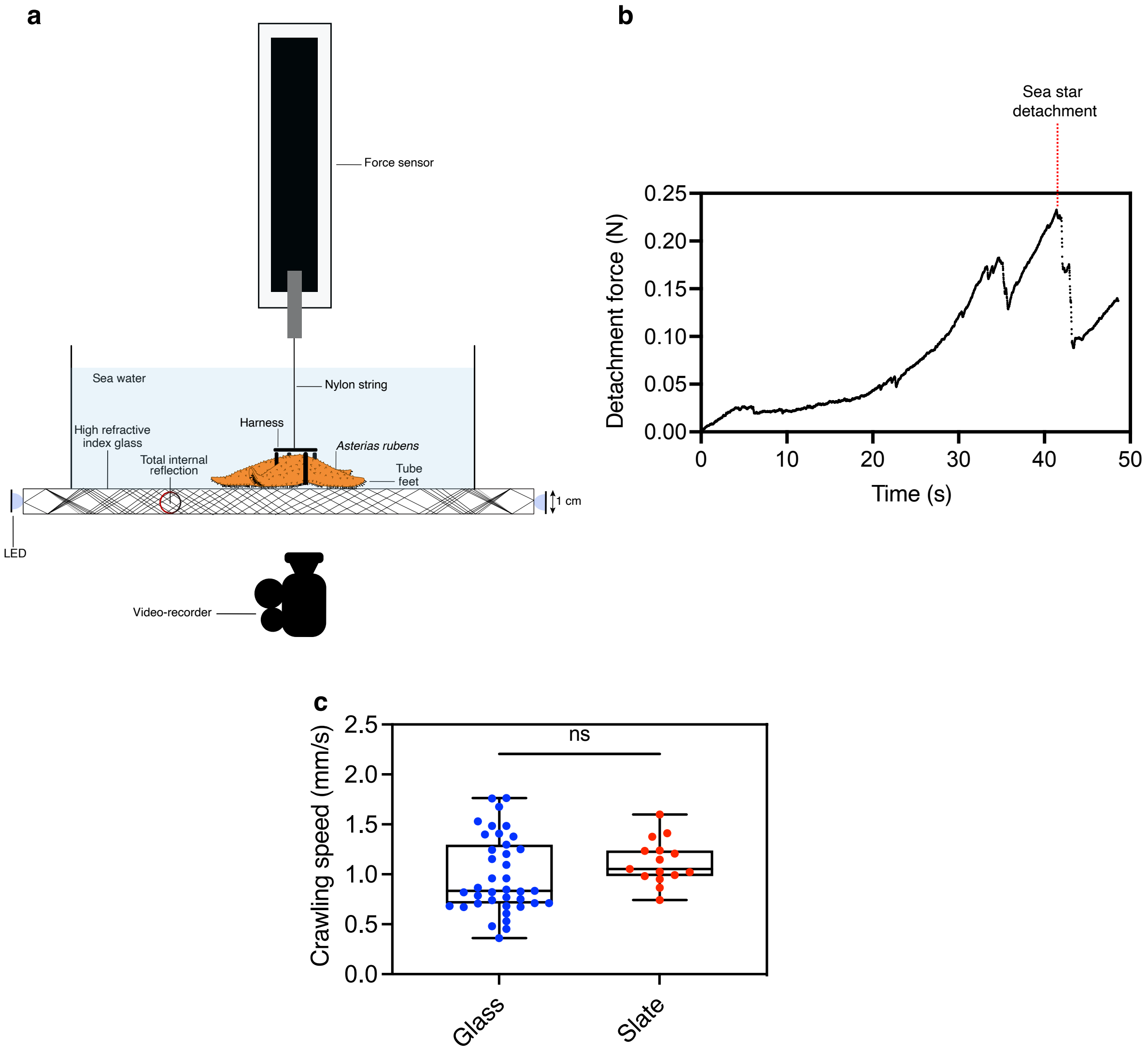


**Supplementary Figure 2 – Glass is a suitable substrate for studying sea star locomotion.** **(a)** Schematic of the experimental setup used to measure detachment force from a glass surface using a force sensor. **(b)** Representative detachment curve showing force as a function of time during removal of one indivudual *Asterias rubens* from the glass substrate. **(c)** Comparison of crawling speed of *A. rubens* on glass ($n$ = 39 *Asterias rubens*) and slate ($n$ = 15 *Asterias rubens*). *p < 0.05; **p < 0.01; ***p < 0.001; ****p < 0.0001; ns = not significant.

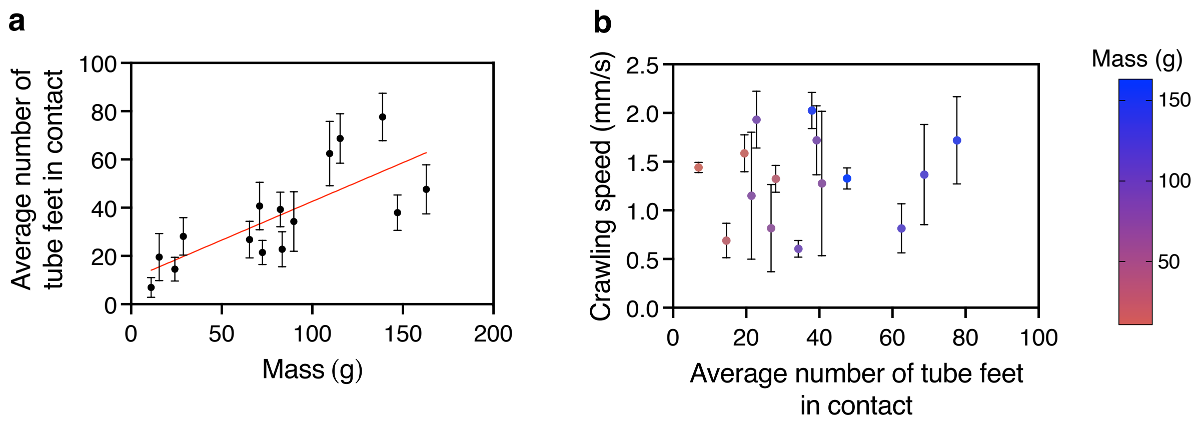


**Supplementary Figure 3 – Crawling speed in *Marthasterias glacialis* is not correlated to the average number of tube feet in contact. (a)** Relationship between the average number of tube feet in contact and body mass in *M. glacialis* ($n$ = 15; $R^2$ = 0.5711, $p$ < 0.0011). **(b)** Relationship between crawling speed and the average number of tube feet in contact during *M. glacialis* locomotion. All data are presented as mean ± standard deviation (s.d.).

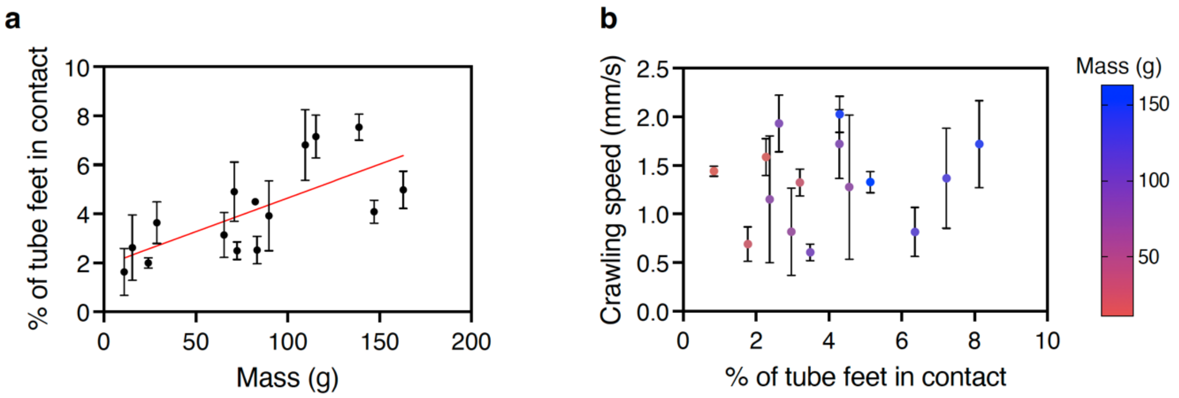


**Supplementary Figure 4 – Percentage of tube feet in contact does not affect crawling speed in *Marthasterias glacialis*. (a)** Relationship between the percentage of tube feet in contact and body mass in *M. glacialis* ($n = 15$; $R^2 = 0.4243$, $p < 0.001$). **(b)** Relationship between crawling speed and the percentage of tube feet in contact ($n = 15$).

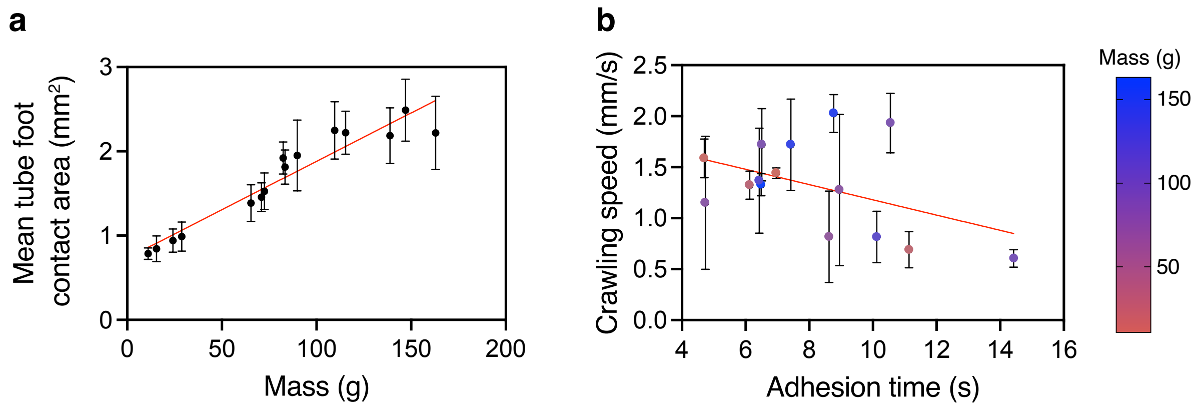


**Supplementary Figure 5 – Tube foot adhesion time supports adaptive and efficient locomotion in *Marthasterias glacialis*. (a)** Relationship between mean tube foot contact area and body mass in *M. glacialis* ($n$ = 15; $R^2$ = 0.7494, $p < 0.0001$). **(b)** Relationship between crawling speed and tube foot adhesion time ($n$ = 15; $R^2$ = 0.1323, $p < 0.0001$). All data are presented as mean ± standard deviation (s.d.).

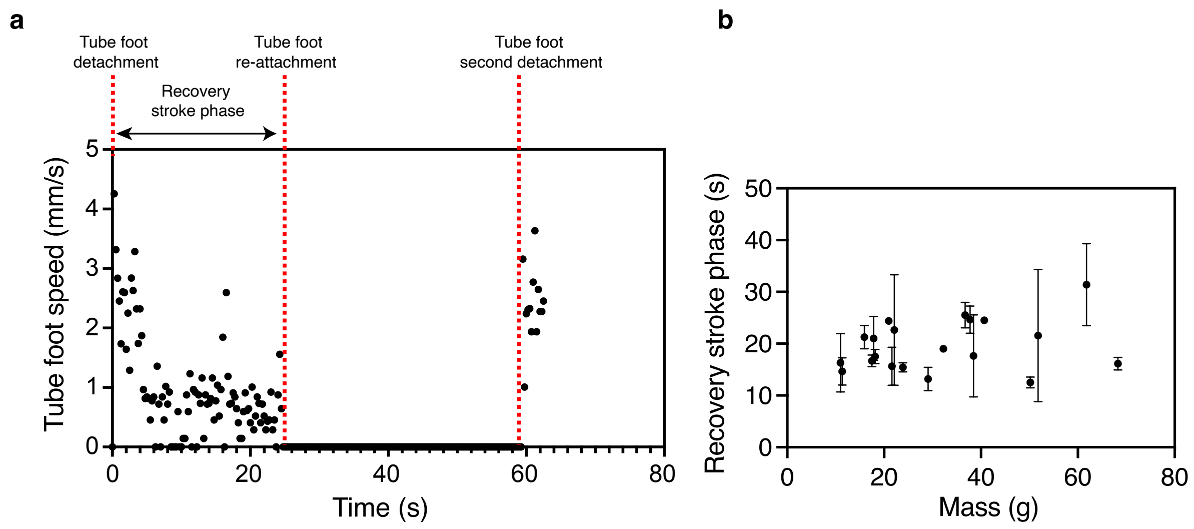


**Supplementary Figure 6 – Tube feet are reused during locomotion in *Asterias rubens*, and the recovery stroke phase is not influenced by the sea star mass.** **(a)** Tube foot speed as a function of time, measured over a 60-second interval. **(b)** Relationship between the duration of the recovery stroke phase and body mass in *Asterias rubens* ($n$ = 20).

**Supplementary Movie 1 –** *Asterias rubens* is vertically detached from a glass surface using pull off force at a constant speed of 1 mm/s using a Zwick Roell force sensor. Frame interval: 0.1 s. Scale bar is 10 mm.

**Supplementary Movie 2** – Side view of *Asterias rubens* arm during locomotion. Tube feet successively attach and detach, enabling force generation for movement. Frame interval is 0.04 s. Scale bar is 5 mm.

**Supplementary Movie 3** – Demonstration of total internal reflection-based system using human fingertips. Bright contact points appear where fingers touch the surface, illustrating the principle of frustrated total internal reflection. Frame interval: 0.1 s. Scale bar: 5 cm.

**Supplementary Movie 4** – Image analysis pipeline developed in Fiji (ImageJ) to detect tube feet in contact during *A. rubens* locomotion. From left to right: raw video, contrast-enhanced video, and thresholded video showing detected tube feet as black dots. Frame interval: 0.2 s. Scale bar: 10 mm.

**Supplementary Movie 5** – Tracking of successive tube foot adhesions during *A. rubens* locomotion. Frame interval: 0.25 s. Scale bar: 10 mm.

**Supplementary Movie 6** – Top view of *A. rubens* locomotion while equipped with a 3D-printed backpack used in perturbation experiments. Frame interval: 0.2 s. Scale bar: 20 mm.

**Supplementary Movie 7** – Simulated sea star locomotion in response to a 25% and 50% increase in initial mass, with no parameter adjustment and all parameters held constant.

**Supplementary Movie 8** – Image analysis method developed in Fiji (ImageJ) for detecting tube feet during *Asterias rubens* inverted locomotion. From left to right: original video of *A. rubens* crawling upside down, followed by the thresholded video showing detected tube feet in contact (black dots). Frame interval: 0.2 s. Scale bar: 20 mm.

**Supplementary Movie 9** – Simulated sea star locomotion under inverted conditions, with no parameter adjustment and all parameters held constant.

### Supplementary Information:
### Tube feet dynamics drive adaptation in sea star locomotion

Amandine Deridoux[1,2], Sina Heydari[3,4], Eva Kanso[3,5], Patrick Flammang[2] and Sylvain Gabriele[1]

[1] Mechanobiology & Biomaterials Group, University of Mons, Research Institute for Biosciences, CIRMAP, Place du Parc, 20 B-7000 Mons, Belgium

[2] Biology of Marine Organisms and Biomimetics Unit, Research Institute for Biosciences, University of Mons, Mons, Belgium

[3] Aerospace and Mechanical Engineering, University of Southern California, Los Angeles, CA 90089, USA

[4] Mechanical Engineering, Santa Clara University, CA 95053, USA

[5] Physics and Astronomy, University of Southern California, Los Angeles, CA 90089, USA

We propose a mathematical model for describing the sea star locomotion driven by the action of hundreds of tube feet. The model was first introduced and developed in [1] and later employed with modifications in [2].

**Sea star biomechanics** We modeled the sea star as a rigid body of length $L$, mass $m$, and submerged weight $W$, whose center of mass is located at $(x, y)$ in the vertical plane. The ventral surface of the sea star is lined with $N$ tube feet, anchored to the body at evenly spaced positions, denoted by the signed distance $d_n$ from the sea star's center of mass, with $d_{n+1} - d_n = d$ and $n \in [1, N]$. We characterized each tube foot by its length $\ell_n$ and inclination angle $\theta_n$ from the vertical $y$-axis. When engaged with the substrate, a tube foot distal end is located at $(x_n, y_n)$. Consequently, the state $(\ell_n, \theta_n)$ of the tube feet, and the location of its distal end $(x_n, y_n)$ are related to the position $(x, y)$ of the sea star body via the algebraic constraints

$$x_n + \ell_n \cos\theta_n = x + d_n, \qquad y_n + \ell_n \sin\theta_n = y. \tag{1}$$

When attached and engaged with the substrate, a tube foot exerts an active force $F_a$ on the sea star's body, induced by activation of muscle tissues lining either the podium or ampula, and experiences a passive restorative force $F_p$ due to the resistance of connective tissues (Fig. 1b). Both active and passive forces act along the length of the tube foot and can either push or pull on the sea star's body [1, 3]. When ampullar muscles are activated, the tube foot extends, applying a pushing force, while when the podium muscles are activated, the tube foot contracts, applying a pulling force. To mathematically model these active forces, we used a piecewise linear force-length constitutive relationship inspired by Hill's muscle model [4, 5] (Fig. 1b,c). When actively pushing, $F_a$ is given by

$$F_a(\ell) = \begin{cases} F_{\max}\dfrac{\ell}{\ell_c}, & \ell < \ell_c, \\ \\ F_{\max}\dfrac{(\ell - \ell_{\max})}{(\ell_c - \ell_{\max})}, & \ell_c < \ell < \ell_{\max}, \\ \\ 0, & \text{otherwise.} \end{cases} \tag{2}$$

where $F_{\max}$ is a scalar denoting the maximum active force that can be generated by the tube feet. When actively pulling, a similar expression with negative active force can be written. For the passive force $F_p$, we used a linear spring $F_p = -k_p(\ell_n - \ell_o)$, where $\ell_o$ is a resting length $\ell_o$ at which $F_p$ vanishes. That is, a tube foot $n$ engaged with the substrate applies a total force $F_n = F_a + F_p$ on the sea star body; When tube foot $n$ detaches from the substrate and enters its recovery stroke, the force $F_n$ goes to zero.

The sea star body moves under the collective action of all tube feet. Applying Newton's second law, we arrive at the equations governing the horizontal and vertical displacements $(x, y)$ of the sea star center of mass,

$$\begin{aligned} -c_x\dot{x} - \sum_n F_n \cos\theta_n &= m\ddot{x}, \\ -W - c_y\dot{y} + \sum_n F_n \sin\theta_n &= m\ddot{y}. \end{aligned} \tag{3}$$

Here, $c_x$, and $c_y$ are internal damping parameters that account for all damping forces, including damping from the tube feet. Eqns. (1) and (3) form a differential-algebraic system of $2 + 2N$ equations for $2+2N$ unknowns $(x, y, l_n, \theta_n)$ provided control rules for when the tube feet should actively apply pushing or pulling forces on the sea star's body and when they should attach and detach from the substrate.

**Non-dimensionalization.** To non-dimensionalize the equations of motion, we use the length of the tube foot $\ell_{\max}$ as our characteristic length scale. The system has two nominal time scales: an inertial time scale $T_g = \sqrt{\ell_{\max}/g}$ obtained from balancing the weight and inertial forces and a relaxation time scale $T_d = c_d/k_p$ obtained from balancing the damping and elastic forces ($c_d\ell_{\max}/T_d \sim k_p\ell_{\max}$). Roughly, $T_d$ is the time it takes for the tube foot to relax to its rest length after being stretched or compressed. Sea stars move slowly and their locomotion is dominated by viscous effects; we thus consider $T_d/T_g \gg 1$, indicating an overdamped regime. Using $\ell_{\max}$ and $T_d$ as our characteristic length and time scales, respectively, we rewrite (3) in non-dimensional form, where the non-dimensional mass $\tilde{m}$ is related to the actual mass $m$ via $\tilde{m} = m/\gamma$, where $\gamma = T_d^2/T_g^2 = (c_d^2/k_p^2)/(L/g) \gg 1$.

**Feedback control at the tube foot level.** In our model, we impose no direct control of the sea star center of mass. Tube feet are controlled based on local adaptive feedback mechanisms. Namely, an attached tube foot senses its own state $(\ell_n, \theta_n)$ and responds as follows: depending on $\theta_n$, the tube foot experiences shear either in the same or in the opposite direction to its horizontal motion, and accordingly, it decides to either push or pull (Fig. 1d). When the tube foot is axially stretched beyond a certain length $l_{\text{detach}}$, it detaches. In the detached state, it applies no force on the sea star body. It stays in the detached state for a random duration $\tau_n$, dictated by a probability of reattachment $P_{\text{reattach}} = \lambda\tau_n$, where $\lambda$ is the rate of reattachment. That is, the longer the duration of the tube foot in the detached state, the higher its probability of reattaching. When a tube foot reattaches, it does so by taking a random step size $\Delta\theta_n$ drawn from a uniform distribution $\Delta\theta_n \sim U(0, \pi/4)$ in the direction of motion of the sea star body, such that, on a flat horizontal terrain, the attachment site $(x_n, y_n)$ satisfies $x_n = x + d_n + y \tan \Delta\theta_n$ and $y_n = 0$.

**Parameter values** In this study, we used measurements in *Asterias rubens* [6, 1] to guide the choice of parameter values in the model. Namely, we set the dimensionless tube foot length to $\ell_{\max} = 1$ and sea star bodylength to $L = 40$, corresponding in dimensional form to about 2 mm tube feet and 8 cm body diameter. We considered $N = 100$ tube feet, and we set a base submerged weight to $W = 2$ and chose the maximal active force to be $F_{\max} = 0.4$. With this choice, 10% of the tube feet exerting on average an active force $F_{\max}/2$ are sufficient to carry the sea star weight, consistently with experimental measurements in *Asterias rubens* [6, 1]. We considered overdamped motion with ratio of relaxation to inertial time scales $\gamma = 50$, and we assumed that a rate of reattachment of $\lambda = 5$.

**Results** We tested numerically the two scenarios that were explored experimentally in the main manuscript: (i) locomotion on a horizontal substrate while increasing the sea star weight, and (ii) locomotion on horizontal and inverted substrate.

In the first set of simulations, we tested three different weights: $W = 2$, $W = 2.5$, and $W = 3$. As the weight increased, we adjusted the detachment length parameter $l_{\text{detatch}}$, from $l_{\text{detatch}} = 0.9$, 0.95 and 1, respectively. For each weight we conducted 25 Monte-Carlo simulations, each for a total time duration of $10T_d$. In each simulation, we calculated the crawling speed, defined as the total distance traveled by the center of mass divided by the total simulation time. We also calculated the attachment fraction, $\tau = \sum_{i=1}^{N} T_i^{\text{attached}}/NT$, defined as the sum over all tube feet of the total time each foot spends in attachment $T_i^{\text{attached}}$, divided by the total simulation $T$ times the number $N$ of tube feet. This gives the average proportion of time that a typical tube foot is attached.

In the second set of simulations, the weight and detachment length were kept fixed at $W = 2$ and $\ell_{\text{detach}} = 0.9$. We conducted 25 random simulations for locomotion on a horizontal substrate and 25 for locomotion an inverted substrate, with no parameter adjustment.

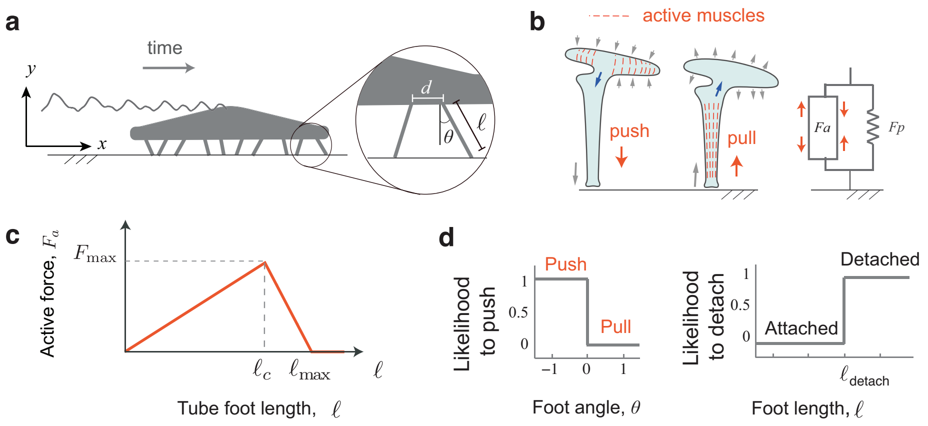


**Supplementary Figure** 1: The mathematical sea star model. **(a)** Locomotion of the sea star model carried by 10 tube feet. The inset shows the length of a foot $l$, its tilt angle $\theta$, and the distance $d$ between two consecutive tube feet. **(b)** Mathematical model of the tube feet consists of a passive linear spring and an active force profile, which can generate either pushing or pulling forces depending on which set of muscles are activated. **(c)** Active force profile inspired by Hill's muscle model. **(d)** Control policies at the tube feet level. The policies indicate the likelihood of transitioning from active pushing to pulling and from attached to detached phase as a function of local sensory cues for each foot. The tube feet transition from detached to attached at length $l_{\text{det}}$.


[3] Ellers O, Ellers KI, Johnson AS, Po T, Heydari S, Kanso E, et al. Soft skeletons transmit force with variable gearing. Journal of Experimental Biology. 2024;227(9):jeb246901.

[4] Hill AV. The heat of shortening and the dynamic constants of muscle. Proceedings of the Royal Society of London Series B-Biological Sciences. 1938;126(843):136-95.

[5] Fung YC, Skalak R. Biomechanics. Mechanical Properties of Living Tissues. Journal of Applied Mechanics. 1982;49:464-5.

[6] Kerkut G. The forces exerted by the tube feet of the starfish during locomotion. Journal of Experimental Biology. 1953;30(4):575-83.
